# Functional repair after ischemic injury through high efficiency *in situ* astrocyte-to-neuron conversion

**DOI:** 10.1101/294967

**Authors:** Yuchen Chen, Ningxin Ma, Zifei Pei, Zheng Wu, Fabricio H. Do-Monte, Pengqian Huang, Emma Yellin, Miranda Chen, Jiuchao Yin, Grace Lee, Angélica Minier-Toribio, Yi Hu, Yuting Bai, Kathryn Lee, Gregory J. Quirk, Gong Chen

**Affiliations:** Department of Biology, Huck Institutes of Life Sciences, Pennsylvania State University, University Park, PA 16802, USA.; Departments of Psychiatry and Anatomy & Neurobiology, University of Puerto Rico School of Medicine, P.O. Box 365067, San Juan, Puerto Rico 00936-5067; Guangdong-Hongkong-Macau Institute of CNS Regeneration, Jinan University, Guangzhou 510632, China.

**Keywords:** NeuroD1, ischemic injury, brain repair, astrocyte-to-neuron conversion, motor function, fear conditioning learning

## Abstract

Mammalian brains have largely lost internal neural regeneration capability except for a few discrete neurogenic niches. After brain injury, the cerebral cortex is especially difficult to repair due to its extremely low rate of adult neurogenesis. Previous studies have converted glial cells into neurons, but the total number of neurons generated is rather limited, casting doubt about its therapeutic potential. Here, we demonstrate that high-efficiency neuroregeneration can be achieved in adult mammalian brains by making use of an engineered AAV Cre-FLEX system to convert a large number of reactive astrocytes into functional neurons. Specifically, using a combination of GFAP::Cre and FLEX-NeuroD1 AAV system, we were able to regenerate enough new neurons from astrocytes to cover about 40% of the neurons lost from an ischemic injury (400 NeuN+ new neurons/mm2), compared to previously reported an average of <1% of cortical neurons (2-8 NeuN+ neurons/mm2) in an ischemic-injured adult mammalian cortex. Importantly, this in situ astrocyte-to-neuron conversion process also improved survival of injured pre-existing neurons, (additional 400 neurons/mm2), leading to a repaired motor cortex with layered cortical structures. Moreover, NeuroD1-converted neurons not only form functional neural circuits but also rescue motor and memory deficits after ischemic injury. Our results establish the proof-of-principle that a highly efficient in situ astrocyte-to-neuron conversion approach provides a novel treatment for neurological disorders that are in need of new neurons.

## INTRODUCTION

Neuronal loss is a major pathological hallmark of brain injury. Regenerating new neurons to replenish lost neurons after an injury is critical for the brain repair. Unfortunately, adult mammalian brains have largely lost active neurogenesis capacity, except a few neurogenic niches such as hippocampus and subventricular zone (SVZ) ^1^. For example, after ischemic injury, the number of new neurons generated through the internal neurogenesis is reported to be less than 1% of total lost neurons in adult mammalian brains ^2–5^. Interestingly, even the neural stem cells (NSCs) originating from the SVZ mostly produce reactive astrocytes, not neurons, after migrating to injured cortical areas ^6, 7^. As an alternative, engrafting external NSCs has been explored for post-stroke treatment ^8–10^, but immunorejection, tumorigenesis, and long-term survival are challenges to this approach ^11, 12^. Therefore, it is urgent to develop new approaches to regenerate a sufficient number of new neurons in order to achieve long-term functional recovery after the brain injury.

We have recently demonstrated the direct conversion of reactive astrocytes into functional neurons by a single transcription factor NeuroD1 in the mouse brain ^13^. Other groups also reported conversion of glial cells into neurons both *in vitro* and *in vivo* ^13–21^. While the *in vivo* glia-to-neuron conversion approach can regenerate new neurons inside mouse brain and spinal cord, it is unclear whether this emerging new technology can generate sufficient numbers of new neurons for therapeutic applications. In the present study, we report that by using an engineered AAV Cre-FLEX system to ectopically express NeuroD1 in reactive astrocytes in a focal ischemic injury model, we are able to regenerate 400 new neurons per mm^2^ in the mouse motor cortex and rebuild the layered cortical structures. Behavioral tests indicate that NeuroD1-treatment significantly rescues both motor deficits and fear memory deficits after the ischemic injury in rodents. Together, our studies demonstrate that NeuroD1-mediated *in situ* cell conversion not only converts astrocytes into functional new neurons but also rebuilds neural circuits and rescues behavioral deficits, establishing a novel approach for functional brain repair following the neural injury.

## RESULTS

### High efficiency of neuroregeneration achieved by AAV Cre-FLEX-NeuroD1 system

We have recently demonstrated that NeuroD1 acts as a master transcription factor to directly convert glial cells into functional neurons inside mouse brains ^13^. However, the total number of neurons generated is rather limited, raising concerns whether this novel *in situ* cell conversion technology has real clinical value^22^. In this study, we used focal ischemic injury as a model system to investigate whether the *in situ* cell conversion approach can achieve functional brain repair. We elicited focal ischemic injury by injecting vasoconstrictive peptide endothelin-1 (ET-1, 1-31) into the motor cortex of FVB mice, producing a significant tissue damage progressively up to 10 weeks following the ischemic injury (Fig. 1a,b) ^23–26^. To convert reactive astrocytes into neurons, we surveyed the injury areas at different time points after ischemic insult and found that astrocytes were not reactive at five days post stroke (dps) (Fig. 1c) but then started to show a significant accumulation of GFAP, a commonly used reactive astrocyte marker, at 10 dps (Fig. 1d). Consistent with our previous report ^13^, injection of retroviruses expressing NeuroD1 at 10 dps resulted in successful conversion of reactive glial cells into NeuN-positive neurons (Fig. 1e). However, the number of neurons was limited due to the fact that retroviruses were only able to express NeuroD1 in the dividing reactive glial cells. To increase the number of neurons, we used adeno-associated virus (AAV) to infect both dividing and non-dividing glial cells. AAV has advantages such as high infection rate and low pathogenicity in humans, and importantly, has been approved by FDA for clinical trials in the treatment of CNS disorders ^27^. To infect astrocytes specifically, we firstly constructed an AAV vector (recombinant serotype AAV9) expressing NeuroD1 under the direct control of a human GFAP promoter (hGFAP::NeuroD1-P2A-GFP). As expected, AAV NeuroD1 successfully converted astrocytes into NeuN-positive neurons (Fig. 1f). While the number of neurons increased significantly using AAV hGFAP::NeuroD1-GFP in comparison with that of retroviruses, the hGFAP promoter can be down-regulated during the astrocyte-to-neuron (AtN) conversion, making this process less complete and less efficient. The reduction of GFP expression under the direct control of GFAP promoter also makes it difficult to distinguish the converted from non-converted neurons. In order to target astrocytes specifically for cell conversion while keeping a high expression level of NeuroD1, we decided to separate the GFAP promoter and NeuroD1 into two different AAV vectors by using the Cre-FLEX (flip-excision) homologous recombination system ^28^. One vector contains a human GFAP promoter to drive the expression of Cre recombinase (hGFAP::Cre) in astrocytes. The second vector contains two pairs of heterotypic, antiparallel loxP-type recombination sites flanking an inverted sequence of NeuroD1-P2A-GFP (or NeuroD1-P2A-mCherry) under the control of a constitutive promoter CAG (CAG::FLEX-NeuroD1-P2A-GFP/mCherry) (Supplementary Fig. 1). The advantage of this AAV Cre-FLEX system is that the Cre expression is controlled by the hGFAP promoter to maintain the specificity towards astrocytes, while NeuroD1 expression is driven by the CAG promoter to avoid being silenced during AtN conversion. Indeed, with this Cre-FLEX system, a strong expression of NeuroD1 in reactive astrocytes was confirmed by immunostaining as early as 4 days post-viral injection (4 dpi) (Fig. 1g, top row). Interestingly, some NeuroD1 AAV-infected astrocytes were clearly captured in a transitional stage toward neurons, showing co-immunostaining of both NeuN and GFAP (Fig. 1g, bottom row). Remarkably, using our AAV Cre-FLEX system, we detected a large number of NeuroD1-converted neurons in the injured mouse cortex (Fig. 1h, i), which was not possible using the other viruses we had tried.

**Figure 1.**
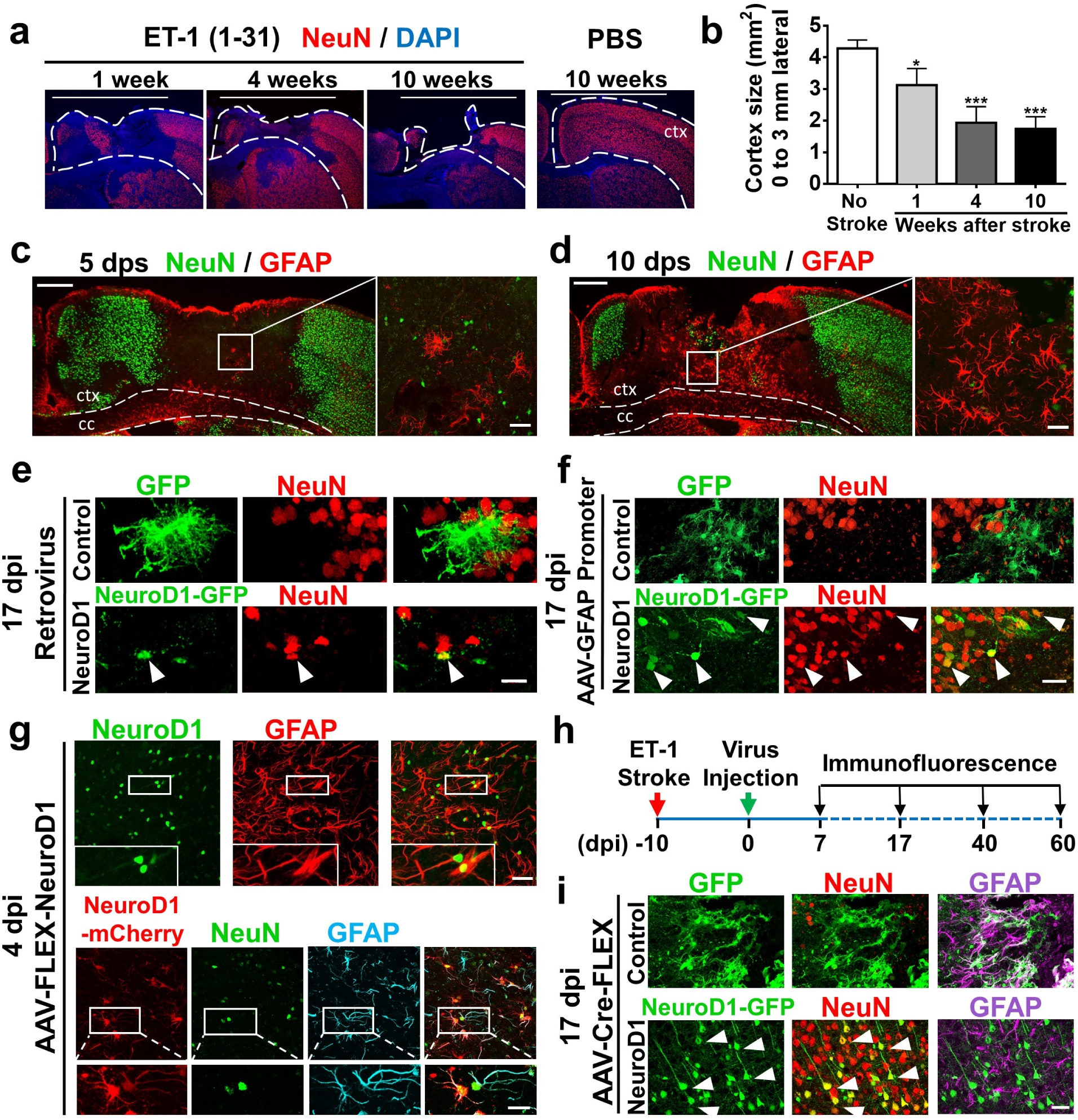
NeuroD1-mediated astrocyte-to-neuron conversion in a focal ischemic injury model. **(a)** Tissue loss caused by focal ischemic injury. Injection of endothelin-1 (ET-1, 1-31) into mouse motor cortex led to a gradual tissue loss in 10 weeks. Dashed lines indicate cortical areas. **(b)** Quantification of the remaining cortical tissue (from the midline to 3 mm lateral area, white bar) at 1, 4 and 10 weeks after ischemic injury in the motor cortex (n = 3 for each time point). **(c, d)** Assessing reactive astrocytes after ischemic injury. Immunostaining of NeuN and GFAP at 5 days (**c**) and 10 days post stroke (dps) (**d**) revealed reactive astrocytes at 10 dps. c.c., corpus callosum. ctx, cortex. **(e)** Retroviruses expressing GFP alone (top row) or NeuroD1-GFP (bottom row), illustrating neuronal conversion by NeuroD1. Viral injection at 10 dps and immunostaining at 17 days post viral injection (dpi). **(f)** Injection of AAV9 expressing GFP alone (hGFAP::GFP, top row) or NeuroD1-GFP (hGFAP::NeuroD1-P2A-GFP, bottom row), illustrating more neurons generated by AAV than retroviruses. **(g)** Capture of the transitional stage from astrocytes (GFAP) to neurons (NeuN) at early time points of NeuroD1 expression (4 dpi). Injection of AAV9 hGFAP::Cre and CAG::FLEX-NeuroD1-P2A-mCherry resulted in significant NeuroD1 expression in GFAP-labeled astrocytes (top row). Interestingly, some NeuroD1-mCherry labeled cells showed both NeuN and GFAP signal (bottom row), suggesting a transition stage from astrocytes to neurons. **(h)** Experimental outline for ischemic injury, AAV injection (Cre-FLEX system), and immunostaining analysis. **(i)** Detection of a large number of NeuroD1-converted neurons using AAV Cre-FLEX system. At 17 dpi, the GFP control group showed many GFAP+ reactive astrocytes (top row, GFAP in purple), whereas the majority of NeuroD1-GFP labeled cells became NeuN-positive (red) neurons (bottom row). Scale bar: **(a)** 3 mm; **(c, d)** Left panel 200 μm, Right panel 40 μm. **(e)** 20 μm. **(f, i)** 40 μm; **(g)** 40 μm (inset 20 μm).

### Neural repair after astrocyte-to-neuron conversion

After achieving high efficiency of neuroregeneration using AAV Cre-FLEX system, we investigated the total number of neurons in NeuroD1-infected areas (Fig. 2a). Compared to the cortex infected with control viruses, NeuroD1 AAV-infected cortex showed many more NeuN+ neurons (red) and less tissue damages (Fig. 2a, left panels in low power). Interestingly, in the NeuroD1 AAV-infected areas, GFAP-labeled astrocytes (purple) persisted and intermingled with NeuroD1-converted neurons, indicating that astrocytes were not depleted by AtN conversion (Fig. 2a, right panels in high power). Surprisingly, after NeuroD1-mediated AtN conversion, we not only detected NeuroD1 positive neurons (green and yellow) but also many non-converted neurons (Fig. 2b, NeuroD1-, red only). Quantitative analysis revealed that ~40% of NeuN+ neurons in the injury areas were NeuroD1-positive and ~60% of neurons were NeuroD1-negative (Fig. 2c; Control group: NeuN+, 27.1 ± 8.1 / 0.1 mm^2^; NeuroD1 group: NeuroD1-/NeuN+, 64.9 ± 7.9 / 0.1 mm^2^; NeuroD1+/NeuN+, 39.9 ± 2.4 / 0.1 mm^2^; n = 3 mice per group). Notably, the non-converted neurons in the NeuroD1-treated areas also more than doubled the total number of neurons in the control GFP-infected areas (Fig. 2c), suggesting that NeuroD1-mediated AtN conversion significantly enhances the survival of pre-existing neurons. Overall, there were more than three times as many NeuN+ neurons in the NeuroD1-infected areas than the controls. Therefore, efficient *in situ* AtN conversion not only regenerated new neurons but also protected injured neurons from further damage from the surrounding toxic environment.

**Figure 2.**
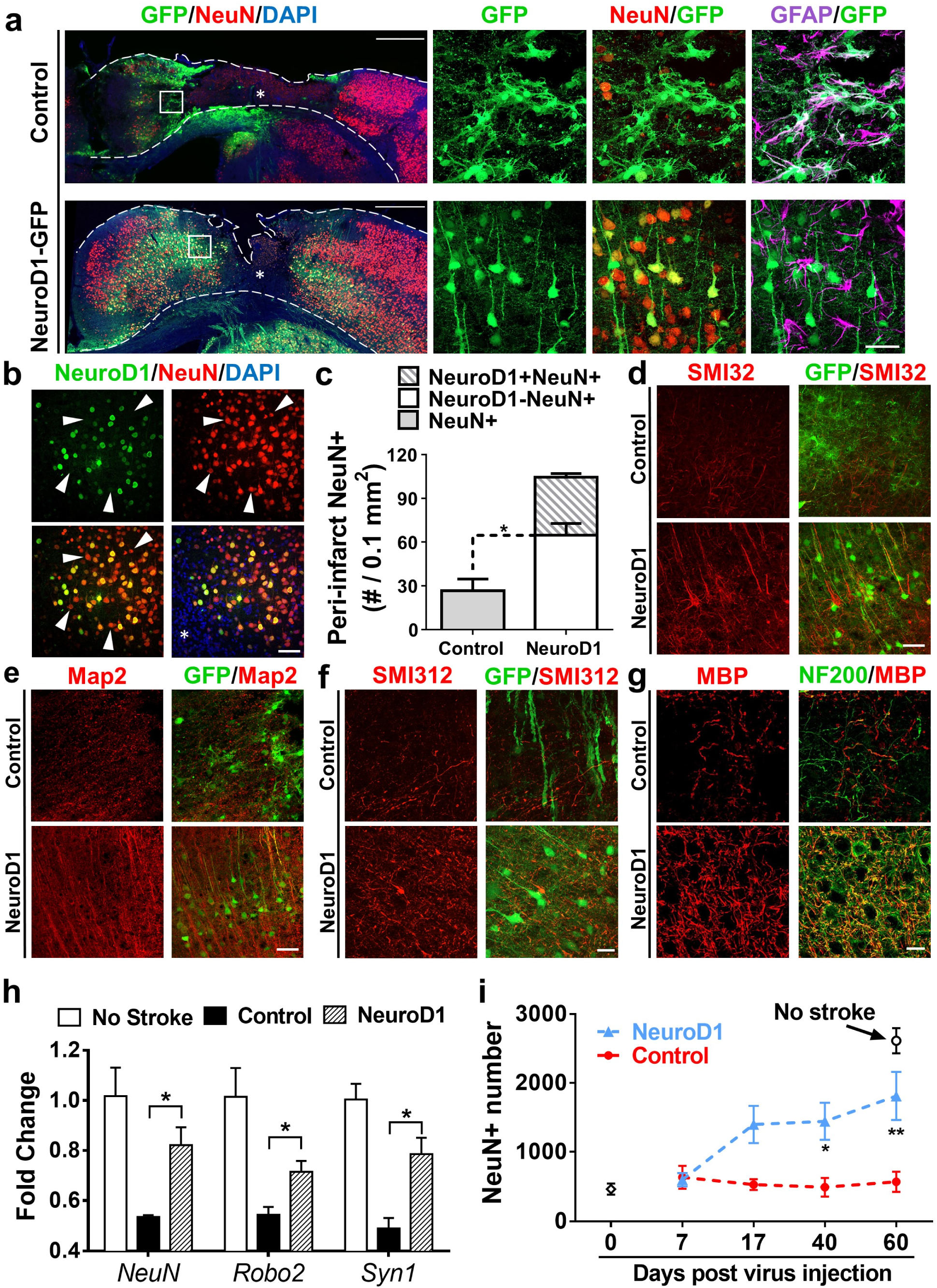
High efficiency of neuroregeneration achieved by NeuroD1-mediated astrocyte-to-neuron conversion in the ischemic injury areas. **(a)** Comparison of the motor cortex (17 dpi) between the control (top row) and NeuroD1 group (bottom row). Left panels showing the overall cortical morphology after ischemic injury and viral treatment, while right panels showing enlarged images of the peri-infarct areas. Note that NeuroD1-infected cells (green) were mostly converted into NeuN+ neurons (yellow), but GFAP+ astrocytes (purple) still persisted in the same areas. **(b)** NeuroD1-converted neurons (green and yellow) were intermingled with non-converted neurons (red, arrowheads) in the injury areas. **(c)** Quantification of total NeuN+ cells in the peri-infarct areas of control group and NeuroD1 group. Note that the number of non-converted neurons in the NeuroD1 group (white bar) more than doubled the number in the control group, suggesting a neuroprotective effect of NeuroD1 conversion. n = 3 mice in each group. * P < 0.05. **(d, e)** Immunostaining of neuronal dendrite markers SMI32 **(d)** and MAP2 **(e)** showing much improved neuronal morphology in the NeuroD1 group (bottom row) compared to the control group (top row). **(f, g)** Immunostaining of axonal marker SMI312 **(f)**, NF200 **(g)** and axon myelination marker MBP **(g)** showed increased axons and axonal myelination in the NeuroD1 group (bottom row) compared to the control group (top row). **(h)** RT-PCR analysis revealed a significant increase of neuronal mRNA level including NeuN, Robo2 and Syn1 after NeuroD1 treatment. * P < 0.05. n = 4 mice each group. **(i)** Quantification of the total number of NeuN+ cells in the motor cortical areas (500-2500 μm lateral from the midline). Note a significant increase of the total number of neurons in the NeuroD1 group by 60 dpi. n = 3 mice in each group. * P < 0.05, ** P < 0.01, Two-way ANOVA followed by Sidak’s multiple comparison tests. Data are represented as mean ± s.e.m. Scale bar: **(a)** 500 μm for left panels; 40 μm for right panels; **(b, d, e)** 40 μm; **(f, g)** 20 μm.

In accordance with a significant increase in the total number of neurons after NeuroD1-treatment, immunostaining of neuronal dendritic markers MAP2 and SMI32 showed clearly aligned dendrites in the NeuroD1 group, compared to the rather disorganized pattern in GFP controls (Fig. 2d,e). Similarly, using axonal marker SMI312 and NF200 along with myelination marker MBP, we found more axons in the NeuroD1 group (Fig. 2f), and these were also highly myelinated (Fig. 2g). Signal intensity and coverage area for these markers are shown in Supplementary Fig. 2. To corroborate with the immunostaining results, we further performed RT-PCR experiments and found that the expression of neuronal genes including *NeuN*, *Robo2*, and *Syn1* all showed a remarkable decrease after the ischemic injury, but significantly rescued by NeuroD1-treatment (Fig. 2h). Moreover, we quantified the total number of NeuN+ neurons within the ischemic areas at different time points following AAV infection. After ischemic injury, the total number of NeuN-positive neurons in the control group was only ~20% of the non-injured brains, indicating that 80% of neurons were lost or injured in our severe focal ischemic injury model. However, in the NeuroD1 group, the NeuN+ neurons showed a significant recovery over the 60 days following viral injection and reached ~70% of the non-injury level (Fig. 2i). Such a significant increase in the total number of cortical neurons after NeuroD1-treatment provides sufficient number of neurons for further functional recovery.

### Reconstitution of motor cortex after neuronal conversion

One pivotal question regarding *in situ* AtN conversion is whether the newly converted neurons can form functional neural circuits. To answer this question, we first examined the cortical structures after ischemic injury with or without NeuroD1-mediated AtN conversion. Over the time course of two months, we observed a gradual cortical tissue loss in the control group injected with the control virus (Fig. 3a, top row), indicating the severe brain injury. In contrast, the cortical tissue was largely preserved in the NeuroD1-treated group without significant atrophy in the motor cortex area (Fig. 3a, bottom row). Notably, after 60 days of NeuroD1 infection, we observed clearly layered structures in the motor cortex of NeuroD1-treated animals (Fig. 3a and 3c, 60 dpi, NeuroD1 images), suggesting reconstitution of cortical circuits after NeuroD1-treatment. Quantitative analysis of the motor cortex revealed that the control group lost 70% of the injured tissue over a 2-month period, whereas the NeuroD1-treated group only lost 20% (Fig. 3b). Besides tissue preservation, we further discovered that the NeuroD1-converted neurons sent their projections globally to many brain regions, including ipsilateral and contralateral areas. For example, a sagittal section of the mouse brain revealed that the NeuroD1-converted neurons projected axons from the motor cortex to the ipsilateral striatum, thalamus, and hypothalamus (Fig. 3c). Coronal section of the motor cortex showed that NeuroD1-converted neurons projected their axons to the contralateral cortical regions through the corpus callosum (Supplementary Fig. S3). Thus, NeuroD1-converted neurons can integrate into global brain circuits.

**Figure 3.**
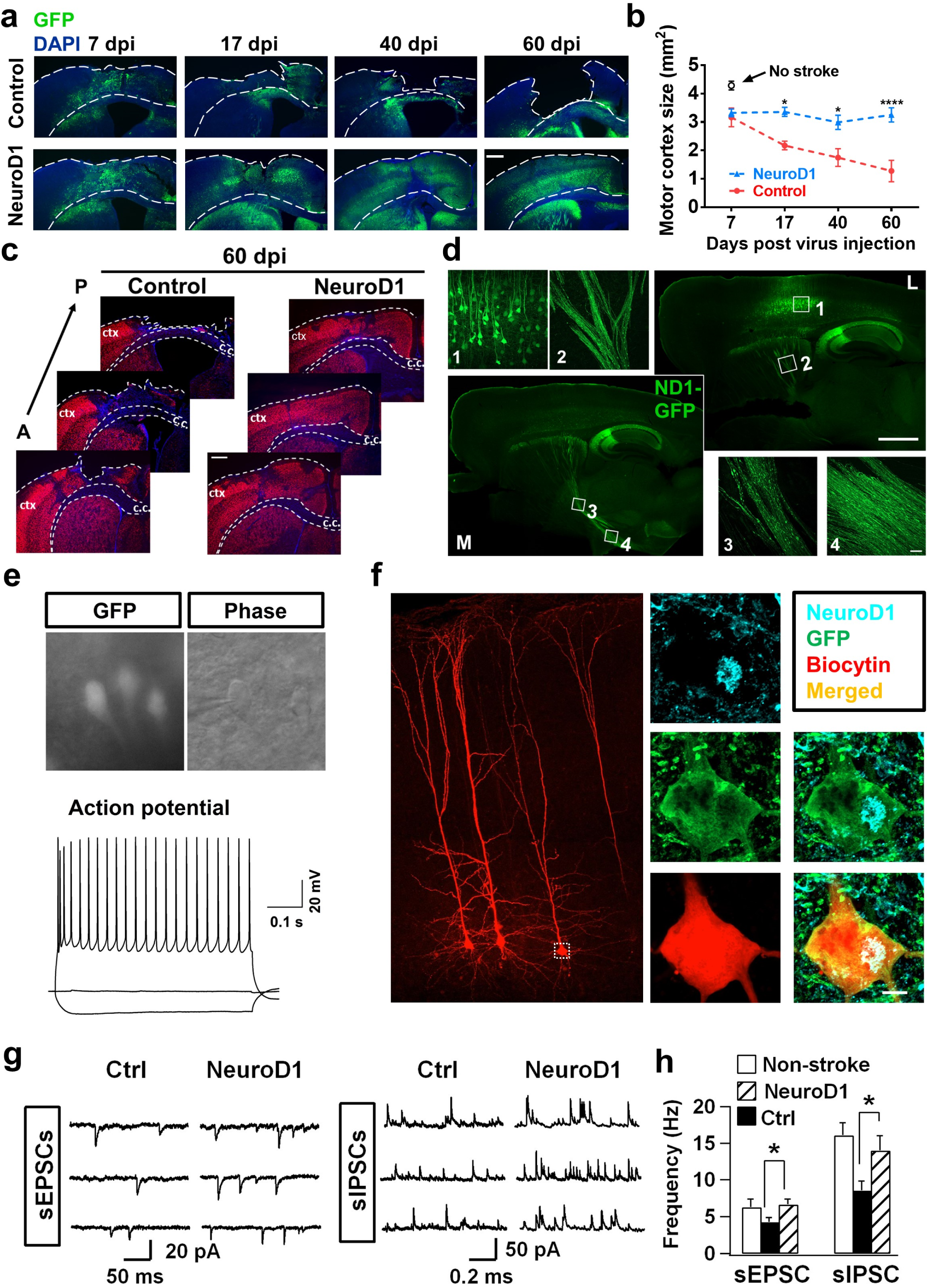
Reconstitution of motor cortex with layered structures after NeuroD1-treatment. **(a)** Low magnification images illustrating gradual tissue loss in the GFP control group (top row), and the rescue by NeuroD1 treatment (bottom row). Note layered structures in NeuroD1 group at 60 dpi. Dashed lines delineate the cortical areas. **(b)** Quantification of the motor cortical areas (from midline to 3 mm lateral) in the control versus NeuroD1 group. * P < 0.05, *** P < 0.001, Two-way ANOVA followed by Sidak’s multiple comparison test. n = 3 mice per group. **(c)** Serial brain sections from anterior (A) to posterior (P) further illustrating severe tissue damage in the control group. ctx, cortex; c.c., corpus callosum. **(d)** Representative images illustrating global axonal projections from NeuroD1-converted neurons. Serial sagittal sections (17 dpi), from medial (M, lower left) to lateral (L, upper right), showing converted neurons in the cortex (inset 1), axonal bundles in the striatum (inset 2), thalamus (inset 3), and hypothalamus (inset 4). **(e)** Brain slice recording on NeuroD1-converted neurons (GFP) detected repetitive action potential firing (60 dpi, n = 22). **(f)** Biocytin (red) infusion in the recording pipette during whole-cell recording followed with post-fix immunostaining illustrating that the recorded neurons were indeed NeuroD1+ (cyan, 60 dpi). **(g)** Representative traces of spontaneous excitatory (sEPSCs) and inhibitory synaptic events (sIPSCs) recorded in NeuroD1-GFP labeled neurons (60 dpi). **(h)** Quantification of the frequency of both sEPSCs and sIPSCs in cortical slices without injury (white bar), or with ischemic injury (black bar, GFP control; striped bar, NeuroD1 group). Note that NeuroD1 group showed significantly higher frequency of both sEPSCs and sIPSCs than the control group (EPSC: control, 4.3 ± 0.6 Hz, n = 22; NeuroD1, 6.7 ± 0.8 Hz, n = 25; p = 0.023, Student’s *t* test) (IPSC: control, 8.6 ± 1.3 Hz, n = 22; NeuroD1, 14.0 ± 2.0 Hz, n = 25; p = 0.032, Student’s *t* test). Scale bars: **(a, c)** 400 μm; **(d)** 1000 μm for sagittal images; 40 μm for inset images; **(f)** 200 μm for the left low magnification image; 5 μm for right enlarged images.

### Synaptic connection after neuronal conversion

After morphological analysis of the structural recovery after *in situ* AtN conversion, we further investigated functional recovery after NeuroD1-mediated AtN conversion. Cortical slice recordings were performed on NeuroD1-converted neurons at 2 months after viral injection (Fig. 3e, top images). Injecting depolarizing currents into the NeuroD1-GFP labeled neurons triggered repetitive action potentials in every recorded GFP+ neuron (n = 22) (Fig. 3e, bottom trace). To confirm that the recorded GFP+ neurons were converted by NeuroD1 rather than pre-existing neurons, we included biocytin in our recording pipettes and performed immunostaining post whole-cell recordings (Fig. 3f). As expected, the GFP+ neurons we recorded (labeled by biocytin, red) were indeed NeuroD1-positive (cyan). Note that the biocytin immunostaining also illuminated the complex dendrites of the NeuroD1-converted neurons after 2 months of conversion (Fig. 3f), supporting their structural integration into the motor cortex after *in situ* AtN conversion. Functionally, we detected robust spontaneous synaptic events, both excitatory and inhibitory, in NeuroD1-converted neurons in the injury sites after 2 months of AtN conversion (Fig. 3g), suggesting that these neurons have functionally integrated into the brain circuits through synaptic connections with other neurons. Quantitative comparison between the GFP control and NeuroD1-GFP groups demonstrated that the frequency of both EPSCs and IPSCs in the NeuroD1 group was significantly higher than the control group (Fig. 3h; n = 22). Together, these data demonstrate that NeuroD1-mediated AtN conversion can reconstitute functional neural circuits after brain injury.

### Functional rescue of motor deficits

With a significant level of neuroregeneration in the injury areas after NeuroD1-mediated AtN conversion, we then investigated whether NeuroD1 treatment can rescue motor deficits caused by ischemic injury in the mouse motor cortex. We analyzed mouse forelimb functions using three behavioral tests—food pellet retrieval, grid walking, and cylinder test—at different time points following viral injection (Fig. 4a). To test for food pellet retrieval, mice were deprived of food before the training and test to increase their motivation for the food pellet. Before ischemic injury, normal animals could be trained to retrieve 5-6 pellets on average out of total 8 pellets in 5 minutes. After ischemic injury, their pellet retrieval capability was severely impaired, dropping to ~1 pellet on average in 5 min (Fig. 4b). Then, the ischemic injured animals with similar motor deficits were assigned into two groups for viral injection: one group injected with GFP viruses alone, and the other injected with NeuroD1-GFP viruses. At 10 days after viral injection, there was no significant difference between the two groups; but after 20 days of viral injection, the NeuroD1 group started to show improvement in food pellet retrieval (Fig. 4b). By 60 days after viral injection, the NeuroD1 group reached ~4 pellets/5 min, whereas the GFP control group could only retrieve <2 pellets/5 min (Fig. 4b). Similarly, for the grid walking test, normal animals before injury had a low rate of foot fault, typically ~5% of total steps within 5 minutes, while walking freely on a grid, but the foot fault rate increased to over 10% after ischemic injury (Fig. 4c). After viral injection, the NeuroD1 group showed consistent improvement after 20 - 60 days of treatment, with the foot fault rate decreasing to ~7%, whereas the GFP control group remained at a high foot fault rate >9% (Fig. 4c). We also performed a cylinder test to assess the forelimb function when animals rise and touch the sidewall of a cylinder. Normal animals typically use both forelimbs to touch and push steadily against the sidewall, with a normal touching rate ~85% (using both forelimbs). After a unilateral ischemic injury in the forelimb motor cortex, the function of the impaired forelimb was significantly weakened and the impaired limb often either failed to touch the sidewall or dragged along the wall after briefly touching (Fig. 4d, normal touching rate dropped to ~35%). After 20 - 60 days of viral infection, the NeuroD1 group showed a significant recovery in touching the sidewall with the injured forelimb (~60%), whereas the majority of the GFP control mice still had a weak forelimb affected by the ischemic injury (Fig. 4d). For all three tests, injection of PBS as a sham control did not result in any behavioral deficits (black line in Fig. 4b-d), and injection of ET-1 (1-31) without viral injection produced similar deficits as the GFP control viral injection (orange line in Fig. 4b-d). Therefore, by using three different motor behavioral tests, we demonstrate that NeuroD1-treatment can rescue motor functional deficits following ischemic injury in the mouse motor cortex.

**Figure 4.**
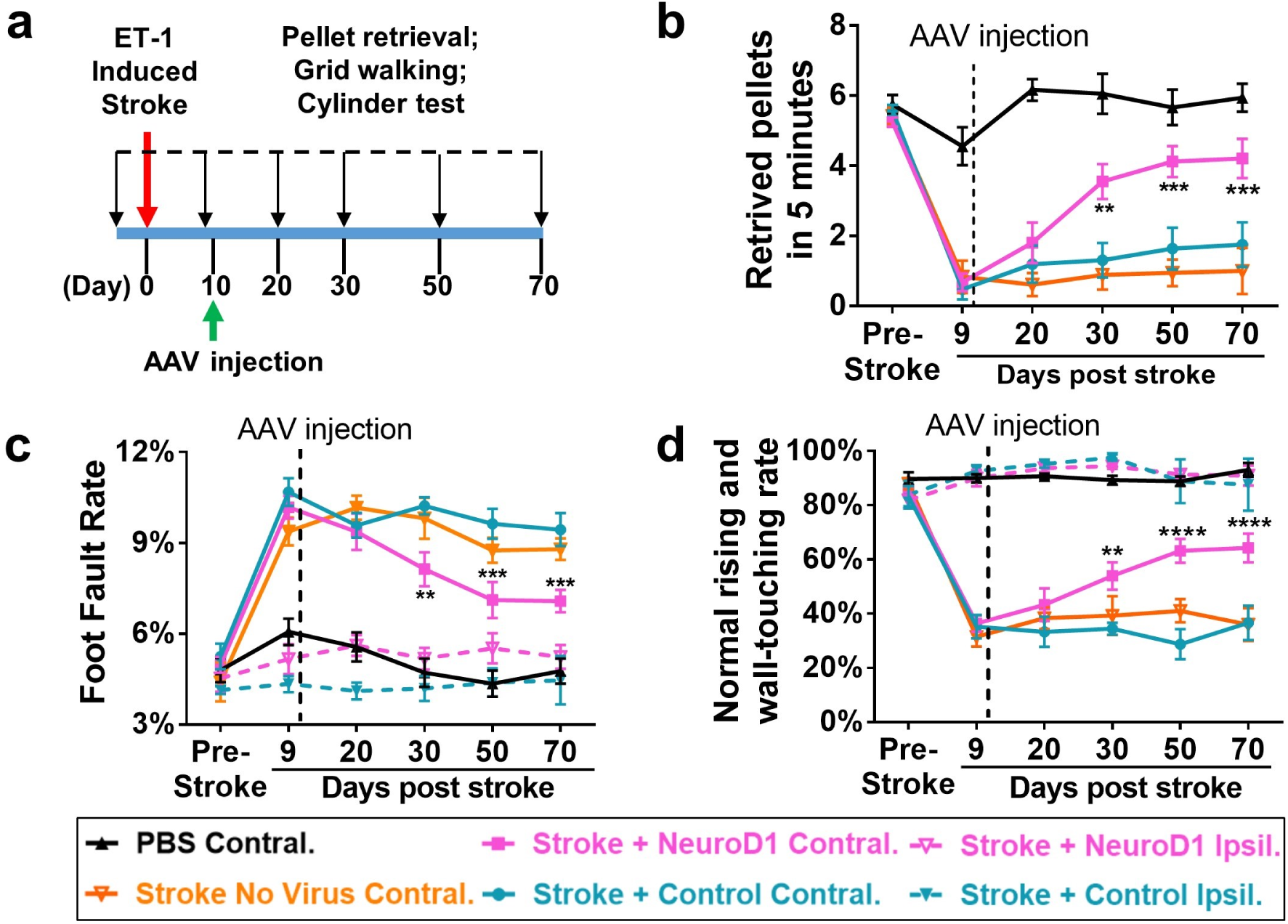
Motor functional improvement after NeuroD1-treatment. **(a)** Experimental design for mouse forelimb motor functional tests. Behavioral tests were conducted before ischemic injury to obtain baseline control, and then 9 dps but one day before viral injection to assess injury-induced functional deficits. AAV were injected at 10 dps, and behavioral tests were further performed at 20, 30, 50, and 70 dps to assess functional recovery. **(b)** Pellet retrieval test. NeuroD1 group (magenta) showed accelerated functional recovery compared to the control group (blue). Ischemic injury at the motor cortex severely impaired the food pellet retrieval capability, dropping from 5-6 pellets/5 min pre-stroke down to 1 pellet/5 min at 9 dps. After NeuroD1 treatment, pellet retrieval ability recovered to 4 pellets/5 min by 60 days post infection. Note that for food pellet retrieval test, only the motor cortex contralateral to the dominant side of forelimb was injured and tested. ** P < 0.01, *** P < 0.001. Two-way ANOVA followed by Tukey multiple comparison test. Data are represented as mean ± s.e.m. n = 12 mice for ET-1 plus control AAV group; n = 11 mice for ET-1 plus NeuroD1 AAV group; n = 6 mice for ET-1 plus no virus group; and n = 6 mice for PBS control group. **(c)** Grid walking test. NeuroD1 group showed lower foot fault rate compared to the control group. Ischemic injury of the motor cortex significantly increased the foot fault rate, which was partially rescued by NeuroD1 treatment. ** P < 0.01, *** P < 0.001. Two-way ANOVA followed by Tukey multiple comparison test. Data are represented as mean ± s.e.m. n = 9 mice for ET-1 plus control AAV contralateral group; n = 11 mice for ET-1 plus NeuroD1 AAV contralateral group; n = 5 mice for ET-1 plus NeuroD1 ipsilateral group; and n = 5 mice for ET-1 plus control ipsilateral group; n = 6 mice for ET-1 plus no virus contralateral group; and n = 6 mice for PBS contralateral group. **(d)** Cylinder test. NeuroD1-treated mice showed considerable recovery of rising and touching the sidewall with both forelimbs compared to the control group. ** P < 0.01, **** P < 0.0001. Two-way ANOVA followed by Tukey multiple comparison test. Data are represented as mean ± s.e.m. n = 9 mice for ET-1 plus control AAV contralateral group; n = 11 mice for ET-1 plus NeuroD1 AAV contralateral group; n = 9 mice for ET-1 plus NeuroD1 ipsilateral group; n = 7 mice for ET-1 plus control ipsilateral group; n = 6 mice for ET-1 plus no virus contralateral group, n = 5 mice for PBS contralateral group.

### Functional rescue of memory deficits

We next used a different animal species (rats instead of mice) and a different behavioral task (cognitive rather than motor) to further test the effect of NeuroD1-mediated AtN conversion following ischemic injury in the amygdala (Fig. 5). It is well established that the associative memory of a conditioned stimulus (tone) and an aversive event (electrical foot shock) is stored in the basolateral nucleus of the amygdala (BLA) ^29, 30^. Lesions of the BLA impair both the acquisition and the subsequent retrieval of an auditory fear memory ^31, 32^. To assess the NeuroD1 effect on cognitive functions, rats were injected with ET-1 into the BLA to produce an ischemic injury and then submitted to auditory fear conditioning three weeks later (Fig. 5a). This interval was based on pilot studies showing a significant impairment in fear acquisition combined with significant neuronal injury approximately 21 days after the ET-1 stroke. The 21-day time point also helped to maximize glial scar formation following ET-1 lesion (Abeysinghe et al., 2014), an important factor to consider when using cell conversion strategies. ET-1 has been previously used to induce ischemic lesions in rat models of stroke ^23, 33^. Accordingly, we found that intra-BLA infusion of ET-1 (1-21) (3 µl/side, 400 pmol) induced a robust lesion in BLA (Supplementary Fig. S4). Behaviorally, rats infused with ET-1 showed a significant reduction in the acquisition of a conditioned freezing response to the tone, in comparison to a saline-infused control group (Fig. 5b, day 21). Reduced freezing was also observed during the retrieval test the following day (Fig. 5b, day 22), suggesting that the lesion induced by ET-1 impaired the formation of an auditory fear memory. One day later, lesioned rats were separated into two groups receiving infusions of either control virus or NeuroD1 virus (3 µl) into the BLA. After three weeks of viral infection, animals were returned to the same box for a fear retrieval test (Fig. 5b, day 45). Rats in the control group continued to show reduced freezing levels, as expected. In contrast, freezing in the NeuroD1 group returned to the levels of the saline/saline group, and was significantly higher than the group receiving the control virus infusion (Fig. 5b, day 45), suggesting a rescue of the memory deficit by NeuroD1 treatment. There was no effect of NeuroD1 on freezing during the pre-tone period (pre-CS, “x” symbol in Fig. 5b), suggesting that NeuroD1 did not induce a general increase in amygdala excitability or non-specific fear. After completion of behavioral tests, we performed immunostaining to confirm viral infection in the BLA. As expected, we found that the majority of NeuroD1-GFP-infected cells were NeuN-positive neurons (Fig. 5c). Together, these results suggest that NeuroD1-treatment can rescue the fear memory deficits induced by an ischemic lesion of BLA.

**Figure 5.**
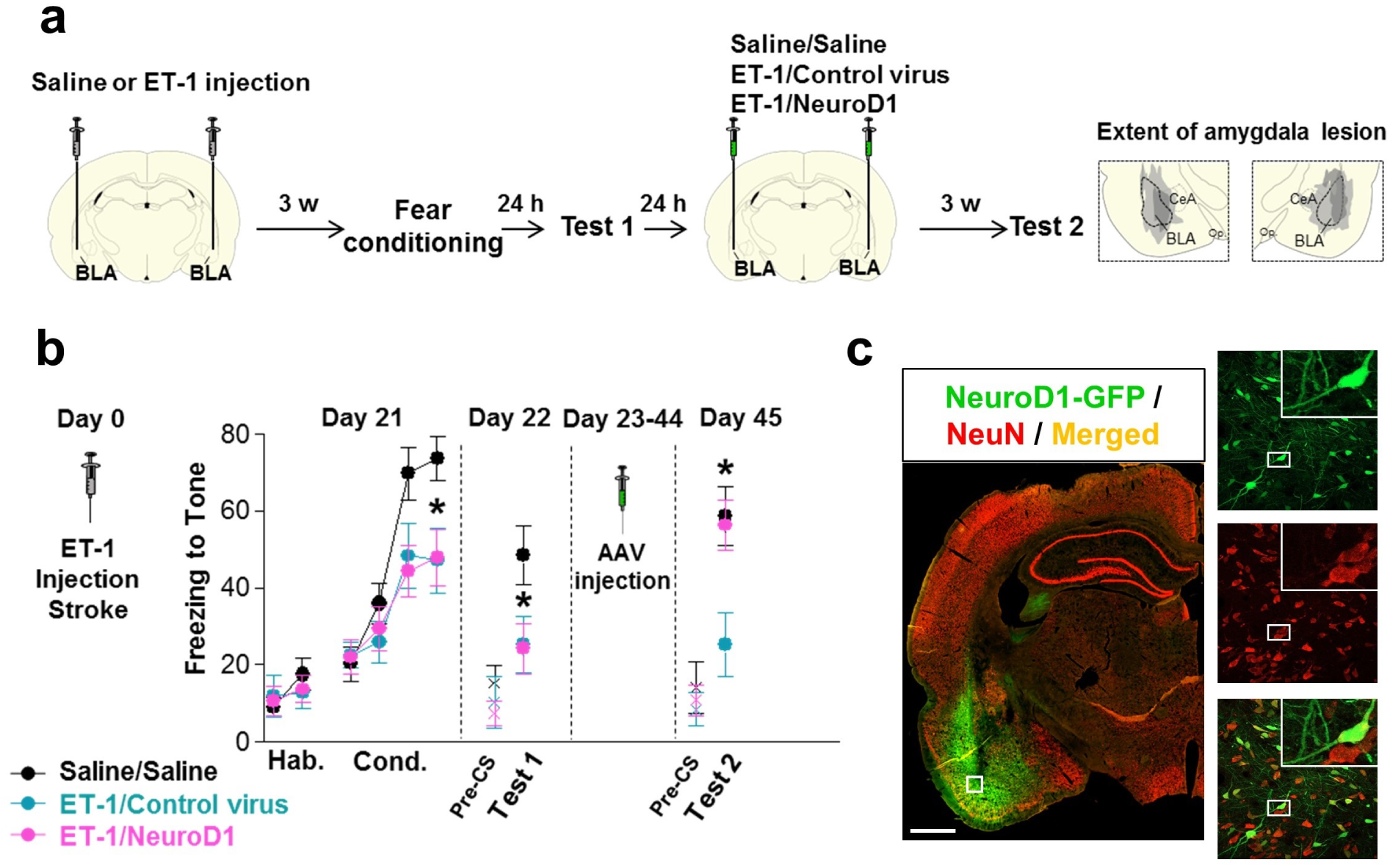
Recovery of fear conditioning memory after NeuroD1 treatment. **(a)** Experimental design of fear conditioning test in rats. ET-1 or saline was injected into the basolateral amygdala (BLA), followed by fear conditioning 3 weeks later. Fear memory tests were performed before viral injection and 3 weeks after viral injection to assess the retention of fear memory. Right two panels illustrate the amygdala lesion induced by the infusion of ET-1. Gray areas represent the minimum (*dark*) and maximum (*light*) spread of the lesion across different anterior-posterior levels of BLA (-2.12, -2.56, and -2.80 from bregma), condensed in one level for illustration. BLA, basolateral nucleus of the amygdala; CeA, central nucleus of the amygdala, op., optical tract. **(b)** ET-1 lesion reduced freezing during fear conditioning (F_(2,31)_ = 3.98, p = 0.02, day 21) at both ET- 1/Control (blue, p = 0.021, n = 10) and ET-1/NeuroD1 (magenta, p = 0.019, n = 14) groups, compared to Saline/Saline group (black, n = 10). Reduced freezing (F_(2,31)_ = 3.45, p = 0.044) was also observed on the next day (day 22) in both ET-1/Control (p= 0.030) and ET-1/NeuroD1 (p = 0.031) groups. Rats were then infused with control virus or NeuroD1 and re-tested 3 weeks later (F(2,31) = 5.86, p = 0.006, day 45). In the ET^-^ 1/NeuroD1 group, freezing returned to the levels of the Saline/Saline group (p= 0.81), and was significantly higher than ET-1/Control group (p = 0.004). “x” denotes baseline pre-tone freezing levels. Hab, habituation; Cond, conditioning; pre-CS, pre-conditioned stimulus. One-way ANOVA followed by Duncan’s post-hoc test. Data are expressed as mean ± s.e.m. in blocks of two trials. * P < 0.05. **(c)** After fear conditioning test, immunostaining of rat brain sections confirmed the injection of NeuroD1-GFP viruses into the BLA (green, left panel), and the NeuroD1-infected cells were mostly NeuN-positive neurons (right panels). Scale bar: 1000 μm.

## DISCUSSION

In this study, we demonstrate that NeuroD1-mediated *in vivo* astrocyte-to-neuron conversion can efficiently regenerate a large number of functional neurons in an ischemic injury model and achieve functional rescue of both motor and cognitive deficits in rodents. The high conversion efficiency is due to the master reprogramming effect of NeuroD1, while the effective neuroregeneration is the result of a combination of the AAV vector and the Cre-FLEX system. One interesting discovery is the persistence of astrocytes at the injury sites after NeuroD1-mediated AtN conversion, indicating that astrocytes would not be depleted after conversion. Another important finding is that besides neuroregeneration from the cell conversion, many injured neurons are preserved after AtN conversion, suggesting a neuroprotective effect when changing reactive astrocytes into functional new neurons. With neuroregeneration and neuroprotection together, we could rescue 70-80% of the lost neurons after ischemic injury and reconstitute the motor cortex with layered structures. This proof-of-principle study demonstrates that a highly efficient neuroregeneration is achievable in adult mammalian brains, opening a new avenue for brain repair using internal glial cells as a source after injury or disease.

### High efficiency of neuroregeneration through *in situ* cell conversion

Most importantly, our NeuroD1-mediated glia-to-neuron conversion can regenerate a large number of new neurons in the stroke areas, averagely 400 NeuN+ cells/mm^2^, which is about 100 times of the internal neurogenesis capability in the adult mouse cortex or striatum after ischemic stroke ^2–5^. Such remarkable neuroregeneration efficacy is also unmatchable by the typically low efficiency of neuroregeneration after stem cell engraftment. We believe that efficient neuroregeneration is critical for functional repair, because small numbers of new neurons either fail to survive in an injury environment or are insufficient to rebuild neural circuits. This may partly explain why modulating endogenous adult neurogenesis through various approaches so far has not yet resulted in an effective therapy for stroke, possibly due to limited neural stem cell niches in the adult brain ^1, 34^. In contrast to modulating endogenous neural stem cells, our *in situ* cell conversion approach makes use of reactive glial cells that are closely associated with neural injury throughout the nervous system for *in situ* regeneration and repair.

As one of the efficient vehicles for gene delivery, AAV has been extensively used in the central nervous system for many purposes, such as gene expression and circuit mapping ^35^. It has also been used as therapeutic vectors for treatments of neurological disorders. AAV has advantages of high infection rate and long-lasting expression with low pathogenicity. For *in situ* AtN conversion, compared to the retroviruses used in the previous study^13^, AAV also has the advantage of infecting both dividing and non-dividing astrocytes, leaving the AtN conversion unrestrained by the number of dividing astrocytes at the viral injection. On the other hand, because AAV can infect both dividing and non-dividing cells, it becomes critical to target glial cells specifically with glia-specific promoters. However, the use of glia-specific promoters also generates a dilemma in which the glial promoter itself become silenced or at least inhibited by the neural transcription factor expressed under the control of the glial promoter. Silencing glial promoter by the neural transcription factor during glia-to-neuron conversion process imposes a negative feedback on the expression level of the neural transcription factor, leading to low conversion efficiency. To overcome this negative feedback effect, we developed a Cre-FLEX system that separates the GFAP promoter from NeuroD1 expression so that NeuroD1 expression will not be inhibited during AtN conversion. This is achieved by targeting astrocytes with astrocyte-specific GFAP::Cre AAV vector but expressing NeuroD1 in a different AAV vector under the control of CAG promoter. After Cre-mediated recombination, NeuroD1 will be highly expressed in astrocytes under a more universal CAG promoter to convert astrocytes efficiently into neurons. With this engineered Cre-FLEX AAV system, we demonstrate here that ectopic expression of NeuroD1 in reactive astrocytes following ischemic injury of mouse motor cortex can regenerate >40% of lost neurons, an unprecedented neuroregeneration efficiency compared to classical pharmacological or genetic interventions, which typically regenerate <1% of lost neurons. For example, previous studies have reported the activation of adult neurogenesis after stroke^5, 36^, but the number of new neurons generated through modulating internal neurogenesis is ~2-8 NeuN^+^ neurons/mm^2 2-5^. When considering the actual neuronal density in the brain, which is ~1000 neurons/mm^2 37^, it may not be too difficult to understand why so many clinical trials on stroke have failed over the past 30 years ^38^. In contrast, our *in vivo* AtN conversion technology makes use of local astrocytes that are surrounding every single neuron to regenerate new neurons wherever neural injury occurs, breaking the bottleneck of classical adult neurogenesis that is limited in certain niches. More importantly, by using the AAV Cre-FLEX system to express NeuroD1 in reactive astrocytes, we now achieve neuroregeneration of ~400 NeuN^+^ neurons/mm^2^, bringing brain repair into a new level with a sufficient number of new neurons to reconstruct the damaged brain structures. With such a high neuroregeneration efficiency, plus additional neuroprotective effect, we demonstrate that the ischemic-injured motor cortex can be gradually reconstituted, and the motor function can be significantly improved.

The use of Cre recombinase might raise concerns regarding its possible toxicity^39 40 41^. However, we designed our AAV system so that Cre is controlled by GFAP promoter, which will be silenced once NeuroD1 is expressed in astrocytes during the early transitional period of astrocyte-to-neuron conversion process. Indeed, this has been confirmed with Cre immunostaining following NeuroD1 infection (Suppl. Figure 5a). In control group (GFAP::Cre + FLEX-GFP), the Cre expression level increased in reactive astrocytes from 4 days to 17 days post infection (dpi), as expected (Suppl. Figure 5a, top row). In NeuroD1 group (GFAP::Cre + FLEX-NeuroD1-GFP), however, Cre was only detected in NeuroD1-GFP positive cells at 4 dpi before conversion but disappeared at 17 dpi, indicating the silence of GFAP promoter once NeuroD1 is expressed (Suppl. Figure 5a, bottom row). Note that in NeuroD1 group, the converted neurons (green) all lacked Cre signal (red), and the remaining Cre signal was not in converted cells.

Another concern is the continuous expression of NeuroD1 in converted neurons after Cre-mediated recombination. In fact, NeuroD1 is an endogenous neural transcription factor that is not only expressed during early brain development but also in the adult neural stem cells as well ^42, 43^. During our immunostaining analysis of NeuroD1 after infection, low level of NeuroD1 expression can be detected in the non-infected pre-existing neurons as well (Suppl. Figure 5b-c). To more directly address whether overexpression of NeuroD1 might impair neuronal functions, we performed brain slice recordings in NeuroD1-GFP converted neurons after 13 months of NeuroD1 infection in mouse brains. The NeuroD1-converted neurons not only survived well after one year, but also displayed repetitive action potentials and synaptic responses (Suppl. Figure 5d), suggesting that the possible side effect of NeuroD1 is minimal, if any.

### Structural support for functional rescue

An exciting finding of this study is that *in situ* AtN conversion can functionally rescue the motor and memory deficits caused by ischemic injury in rodent models. This functional rescue is based on structural changes including both cellular conversion and circuit rebuilding after NeuroD1-treatment. Importantly, we have observed a mixture of NeuroD1-converted neurons together with non-converted neurons in the injury areas after NeuroD1-treatment, suggesting that the repaired brain circuitry is an integration of both newly generated and pre-existing neurons. The fact that NeuroD1-treatment can rescue over 70% of neurons in ischemic injured motor cortex provides necessary structural basis for motor functional recovery. Similarly, a reversal of the memory deficit induced by an amygdala lesion suggests a restoration of the neural circuits in the BLA following NeuroD1 treatment.

### Perspective on therapeutic potential

The molecular and cellular neuroscience research over the past several decades has gained much insight into the brain structure and function as well as the pathological changes after brain injury. However, therapeutic treatments for effective brain repair has significantly lagged behind. In fact, most of the clinical trials on CNS disorders such as stroke and Alzheimer’s disease have largely failed over the recent years ^38, 44^. One of the most important reasons is that previous clinical trials cannot regenerate sufficient number of functional neurons to replace the lost neurons after injury. Because neurons are the fundamental building blocks of brain circuits, it is pivotal to have sufficient number of functional new neurons in order to rebuild the impaired neural networks after injury or disease. For example, if one million neurons die after brain injury, simply generating 10,000 neurons may not be sufficient to achieve significant functional repair. It might require regenerating 100,000 or 200,000 new neurons to start really meaningful neural repair. In other words, it might need regenerating at least 10% to 50% of the lost neurons in order to reconstruct the brain circuits that are disrupted by neural injury. The precise threshold in terms of the percentage of new neurons regenerated, whether 10% or 50% of the lost neurons, may vary according to the type of injury or the location in the CNS. Nevertheless, without sufficient number of new neurons added to the injured neural circuitry, it will be difficult to fully restore the lost brain functions solely relying upon the internal compensation mechanisms. Similarly, any serious neurodegenerative disorders such as Alzheimer’s disease and Parkinson’s disease may also require an effective neuroregeneration approach to reverse the disease symptoms ^45^.

Different from previous pharmacological modulation or stem cell transplantation, our *in vivo* AtN conversion approach makes use of the enormous reservoir of internal glial cells in the CNS to regenerate new neurons. Because glial cells are widely distributed throughout the brain and spinal cord, and more importantly, glial cells can divide to regenerate themselves, the resource for internal neural conversion is enormous. In theory, it is possible to regenerate new neurons anywhere and anytime in the brain so long as the reactive glial cells are surviving the injury. However, if a neural injury is so severe that most glial cells have died, then it may require injection of new cell source such as stem cells and immature neurons, or even glial cells, for neural repair.

With the successful development of the highly efficient neuroregeneration approach that can regenerate 40% of the lost neurons and protect additional 40% of injured neurons, we predict that our NeuroD1-mediated AtN conversion technology is one step closer toward a possible gene therapy for brain repair. For example, if a patient is diagnosed with a focal stroke or focal brain injury under MRI scanning, NeuroD1 AAV particles can be locally injected into the injury areas to regenerate new neurons and restore neuronal functions. From a broader perspective, such *in situ* cell conversion therapy may be applied for tissue repair not only in CNS^18^, but also in a variety of organs such as pancreas, heart, and liver ^46–51^. Importantly, local injection of a small amount of AAV particles may be a preferred route over systemic administration of a large amount of AAV particles, given recent success of local AAV injection in the retina^52^ but a disastrous outcome of intravenous AAV injection^53^.

## ACKNOWLEDGEMENT

This work was supported by grants from National Institutes of Health (AG045656) and Alzheimer’s Association (ZEN-15-321972) to G.C. It was also supported by Charles H. Smith Endowment Fund for Brain Repair and Verne M. Willaman Endowment Fund from the Pennsylvania State University to G.C. The rat fear conditioning test was supported by NIMH grant K99-MH-105549 to F.H.D-M.; NIMH grants R37-MH058883 and P-50-MH-086400 to G.J.Q, and a grant from the University of Puerto Rico President’s Office to G.J.Q. We would like to thank Matthew Keefe and Joseph Gyekis for careful proof reading of our manuscript. We would also like to thank all the Chen lab members for rigorous discussing throughout this project over the past 5 years.

